# Activity subspaces in medial prefrontal cortex distinguish states of the world

**DOI:** 10.1101/668962

**Authors:** Silvia Maggi, Mark D. Humphries

## Abstract

Medial prefrontal cortex (mPfC) activity represents information about the state of the world, including present behaviour, such as decisions, and the immediate past, such as short-term memory. Unknown is whether information about different states of the world are represented in the same mPfC neural population and, if so, how they are kept distinct. To address this, we analyse here mPfC population activity of rats learning rules in a Y-maze, with self-initiated choice trials to an arm-end followed by a self-paced return during the inter-trial interval (ITI). We find that trial and ITI population activity from the same population fall into different low-dimensional subspaces. These subspaces encode different states of the world: multiple features of the task can be decoded from both trial and ITI activity, but the decoding axes for the same feature are roughly orthogonal between the two task phases, and the decodings are predominantly of features of the present during the trial but features of the preceding trial during the ITI. These subspace distinctions are carried forward into sleep, where population activity is preferentially reactivated in post-training sleep, but differently for activity from the trial and ITI subspaces. Our results suggest that the problem of interference when representing different states of the world is solved in mPfC by population activity occupying different subspaces for the world states, which can be independently decoded by downstream targets and independently addressed by upstream inputs.

**Significance statement:** Activity in the medial prefrontal cortex plays a roles in representing the current and past states of the world. We show that during a maze task the activity of a single population in medial prefrontal cortex represents at least two different states of the world. These representations were sequential and sufficiently distinct that a downstream population could separately read out either state from that activity. Moreover, the activity representing different states is differently reactivated in sleep. Different world states can thus be represented in the same medial prefrontal cortex population, but in such a way that prevents potentially catastrophic interference between them.

## Introduction

The medial prefrontal cortex (mPfC) plays key roles in adaptive behaviour, including reshaping behaviour in response to changes in a dynamic environment (Euston et al., 2012) and in response to errors in performance (Narayanan and Laubach, 2008; Laubach et al., 2015). Damage to mPfC prevents shifting behavioural strategies when the environment changes (Laskowski et al., 2016; Guise and Shapiro, 2017). Single neurons in mPfC shift the timing of spikes relative to hippocampal theta rhythms just before acquiring a new action-outcome rule (Benchenane et al., 2010). And multiple labs have reported that global shifts in mPfC population activity precede switching between behavioural strategies (Rich and Shapiro, 2009; Durstewitz et al., 2010; Karlsson et al., 2012; Powell and Redish, 2016) and the extinction of learnt associations (Russo et al., 2021).

Adapting behaviour depends on knowledge of both the past and the present state of the world. Deep lines of research have established that mPfC activity represents information about both. The memory of the immediate past is maintained by mPfC activity in tasks requiring explicit use of working memory (Baeg et al., 2003; Fujisawa et al., 2008; Spellman et al., 2015). The use of such memory is seen in both the impairment arising from mPfC lesions (Rich and Shapiro, 2007; Young and Shapiro, 2009; Laskowski et al., 2016), and the role of mPfC in error monitoring (Laubach et al., 2015). Representations of stimuli and features happening in the present have been reported in a variety of decision-making tasks throughout PfC (Averbeck et al., 2006; Rigotti et al., 2013; Hanks et al., 2015; Siegel et al., 2015), and specifically within rodent mPfC (Sul et al., 2010; Ito et al., 2015; Guise and Shapiro, 2017).

Little is known though about how mPfC activity represents multiple states of the world. Prior studies have shown that past and upcoming choices can both modulate activity of neurons in the same mPfC population (for example Baeg et al., 2003; Ito et al., 2015), but none have compared how different states of the world are represented. Thus important questions remain: if and how different world states are encoded in the same mPfC population, and how those representations are kept distinct.

To address these questions, we reanalyse here mPfC population activity from rats learning new rules on a Y-maze (Peyrache et al., 2009). This task had distinct trial and inter-trial interval phases, and we have previously shown that task features of the preceding trial can be decoded from the population activity in the inter-trial interval (Maggi et al., 2018), showing that mPfC activity in this task depends on the state of the world. We can thus address our key questions here by asking if population activity in the trials also represents the state of the same task features and, if so, how that representation is kept distinct between the trial and inter-trial interval phases.

We find that the mPfC population activity occupies different subspaces between trials and inter-trial intervals, providing a basis for separately representing at least these two distinct states of the world. Consistent with representing world states, task features could be decoded from activity in both the trial and inter-trial interval phases, but were strongly distinct: decoding was of the present features in the trial and predominately of features of the preceding trial during the inter-trial interval. Decoding axes were, or close to, orthogonal between the trials and inter-trial intervals, showing the subspaces supported distinct encodings. Further consistent with their occupying different subspaces, population activity of the trials and inter-trial intervals preferentially reactivated in post-training sleep in different ways: preferential reactivation of trial activity uniquely occurred after learning and correlated with performance during training. Our results thus suggest representing different world states using independently decodable axes within a mPfC population could prevent interference between them, allowing them to be separately accessed by both downstream and upstream populations.

## Materials and Methods

### Task description and electrophysiological data

All data in this study comes from previously published data (Peyrache et al., 2009). Full details of training, spike-sorting, and histology can be found in (Peyrache et al., 2009). The experiments were carried out in accordance with institutional (CNRS Comité Opérationnel pour l’Ethique dans les Sciences de la Vie) and international (US National Institute of Health guidelines) standards and legal regulations (Certificate no. 7186, French Ministeère de l’Agriculture et de la Pêche) regarding the use and care of animals.

Four male Long-Evans rats were implanted with tetrodes in the medial wall of pre-frontal cortex, covering the prelimbic and infralimbic regions, and trained on a Y-maze task (Figure 1a). During each session, neural activity was recorded for 20-30 minutes of sleep or rest epoch before the training epoch, in which rats worked at the task for 20-40 minutes. After that, another 20-30 minutes of sleep or rest epoch recording followed. During the sleep epochs, intervals of slow-wave sleep were identified offline from the local field potential (details in Peyrache et al., 2009; Benchenane et al., 2010).

**Figure 1.**
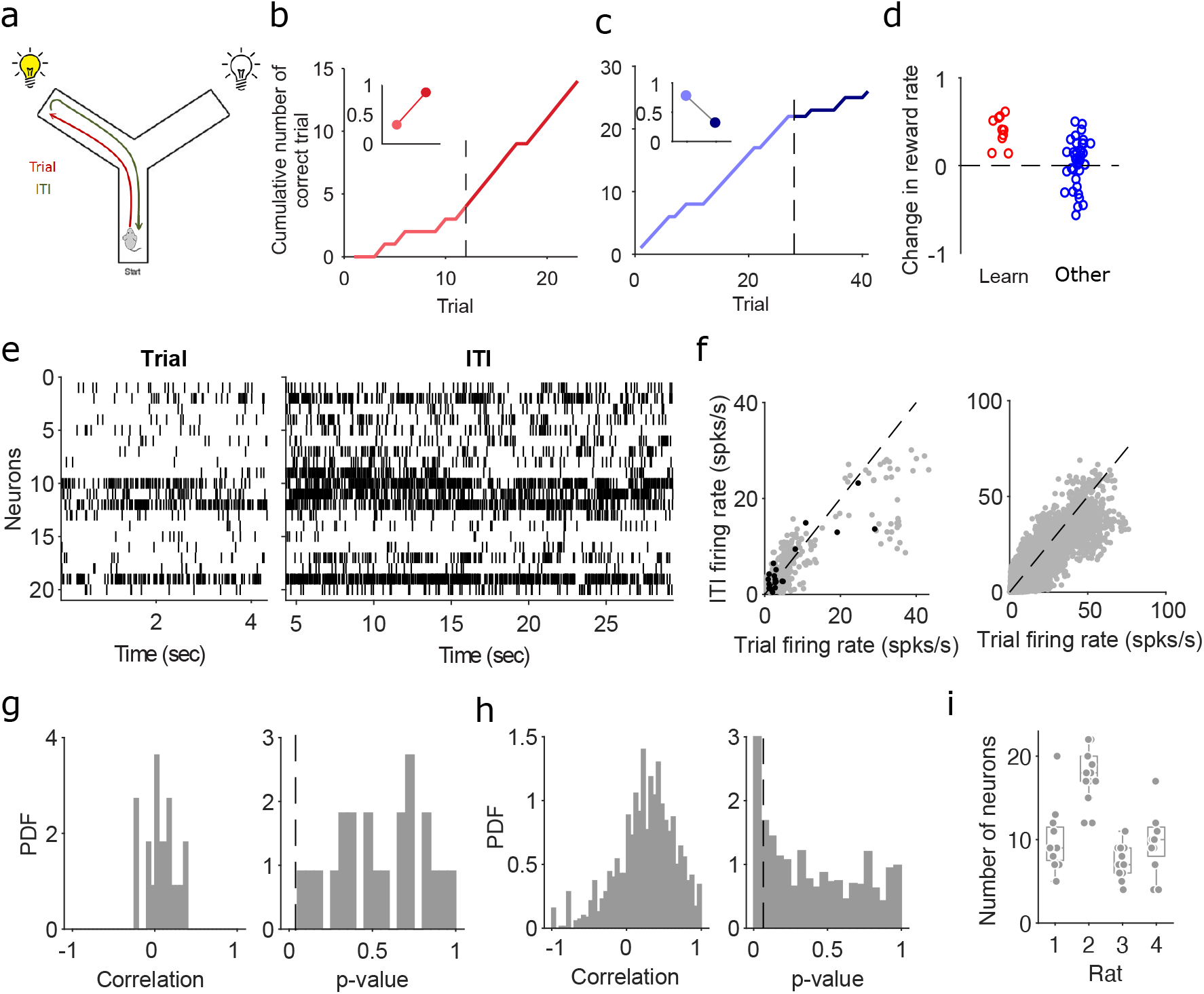
Rule learning and neural activity on the Y-maze task. (**a**) Schematic of the Y-maze task, showing a rat at the start position. A trial is the period from the start position to the end of the chosen arm; the inter-trial interval is the return from the arm end to the start position. On each trial one arm-end was lit, chosen in a pseudo-random order, irrespective of whether it was relevant to the currently enforced rule. Across sessions, animals were asked to learn one of 4 rules in the sequence: go to the right arm, go to the lit arm, go to the left arm, go to the dark arm. Rules switched after 10 correct choices (or 11 out of 12). (**b**) Example reward curve from a learning session, plotting the cumulative number of correct trials. Black dashed line identifies the learning trial as the first of three consecutive correct trials followed by at least 80% correct trials. Inset: reward rates before (light red) and after (dark red) the learning trial. Reward rates were given by the slope of linear regressions fitted to the reward curve before and after the learning trial. (**c**) Example reward curve from an Other session, one of 8 in which the rule switched; the black dashed line identifies the rule change trial. Inset: reward rates before and after the rule change. (**d**) Change in reward rate during all learning sessions (red) or Other sessions (blue). (**e**) Raster plots of spiking activity in the medial prefrontal cortex during a single trial and the following inter-trial interval (ITI). (**f**) Neuron firing rates in each trial and in the following inter-trial interval. Left: an example session, black dots are the data in panel e. Right: all sessions. (**g**) For the same example session, the distribution of Spearman’s rank coefficients between the population vectors of firing rates in the trial and inter-trial interval (left), and the corresponding p-values (right); *p* = 0.05 is indicated by the dashed line. (**h**) as panel g, for all sessions. (**i**) The number of neurons in populations analysed here, by rat. Each symbol is a session.

The Y-maze had symmetrical arms, 85 cm long, 8 cm wide, and separated by 120 degrees, connected to a central circular platform (denoted as the choice point throughout). The two choice arms had a light at the end, one of which was lit during each trial in a pseudo-random sequence. Rats self-initiated a trial by leaving the beginning of the start arm. A trial finished when the rat reached the end of the chosen goal arm. If the chosen arm was correct according to the current rule, the rat was rewarded with drops of flavoured milk. As soon as the animal reached the end of the chosen arm an inter-trial interval started and lasted until the rat completed its self-paced return to the beginning of the start arm. The central platform was raised once the rat passed it to prevent backtracking along the choice arms. The light was extinguished during the return journey – unfortunately from the data available to us it is not clear exactly when (Francesco Battaglia, personal communication).

Each rat was exposed to the task completely naïve and had to learn each rule by trial- and-error. The rules were presented in sequence: go to the right arm; go to the cued arm; go to the left arm; go to the uncued arm. A rule was switched to the next in the sequence when the animal had achieved 10 correct trials in a row, or 11 out of 12. Across the four rats, there were 8 rule switches in total.

The recording sessions taken from the study of Peyrache and colleagues (Peyrache et al., 2009) were 53 in total. Each of the four rats learnt at least two rules, and they respectively contributed 14, 14, 11, and 14 sessions. We used 49 of these sessions for our analysis, of between 7 and 51 trials each. One session was omitted for missing position data, one for consistent choice of the right arm (in a dark arm rule) preventing decoder analyses (see below), and one for missing spike data in a few trials. An additional session was excluded for having only two neurons firing in all trials. Tetrode recordings were spike-sorted within each recording session. Spikes were recorded with a resolution of 0.1 ms. Simultaneous tracking of the rat’s position was recorded at 30 Hz.

### Testing for separable population activity

We evaluated the difference between population activity in the trial and inter-trial intervals of a session by quantifying their separability in a low dimensional space. For consistency with our previous work, for each session we selected the *N* active neurons that fired at least one spike on each trial (Figure 1e), allowing us to directly compare the decoding results obtained here (see below) to those in (Maggi et al., 2018): the populations thus ranged between 4 and 22 neurons (Figure 1i).

We used principal components analysis (PCA) to project the population vectors of a session onto a common set of dimensions. For each session, we constructed a *N*-length vector of neuron firing rates in each trial **r**_*t*_, resulting in the set of population firing rate vectors {**r**_*t*_(1),…, **r**_*t*_(*T*)} across the *T* trials of a session. We then constructed the data matrix **X** from the firing rate vectors of the population, by concatenating trials and intertrial intervals in their temporal order {**r**_*t*_(1), **r**_*I*_(1),…, **r**_*t*_(*T*), **r**_*I*_(*T*)}^*T*^ across the *T* trials of a session; the resulting matrix thus had dimensions of 2*T* rows and *N* (neurons) columns. Applying PCA to **X**, we projected the firing rate vectors on to the top *d* principal axes (the eigenvectors of **X′X**) to create the top *d* principal components. For each set of *d* components, we quantified the separation between the projected trial and inter-trial interval population vectors using a linear classifier (support vector machine), and report the proportion of misclassified vectors. We repeated this for between *d* = 1 and *d* = 4 axes for each session.

### Linear decoding of task features

To predict which task feature was encoded in mPfC population activity we trained and tested a range of linear decoders (Hastie et al., 2009). In the text we report the results obtained using a logistic regression classifier, but for robustness we also tested three other decoders – linear discriminant analysis, linear support vector machines, and a nearest neighbours classifier – and found similar results. The full details of the decoding analysis can be found in Maggi et al. (2018).

Each trial’s task information was binary labelled for three features: outcome (labels: 0, 1), the direction of the chosen arm (labels: left, right), and the arm position of the light cue (labels: left, right). We used leave-one-out cross-validation to decode each feature from population activity, holding out the *i*th trial’s vector **r**_*t*_(*i*), training the classifier on the *N* – 1 remaining trial vectors, and then using the resulting weight vector to predict the feature’s label for the held-out trial. We quantified the accuracy of the decoder as the proportion of correctly predicted labels over all *T* held out trials of a session. The same approach was used for the inter-trial intervals, by constructing the set of firing rate vectors {**r**_*I*_(1),…, **r**_*I*_(*T*)} across the *T* inter-trial intervals.

For decoding at different positions in the maze, we first linearised the maze, divided it into five equally-sized sections, and then computed the *N*-length firing rate vector of the population for each position *p*, **r**^p^_*t*_ in the trial. For each trial *t* = 1,…,*T* of a session and each section of the maze *p* = 1,…, 5, the set of population firing rate vectors {**r**^p^_*t*_(1),…,**r**^p^_*t*_(*T*)} was used to train the cross-validated decoder. The same approach was used for the position-dependent vectors {**r**^p^_*I*_(1),…, **r**^p^_*I*_(*T*)} in the inter-trial intervals of a session.

For each rat and each session, the distribution of outcomes and arm choices depended on the rats’ performance, which could differ from 50%. Therefore, we trained and cross-validated the same classifier on the same data-sets, but shuffling the labels of the task features across trials. In this way we obtained the accuracy of detecting the correct labels by chance. We repeated the shuffling and fitting 50 times and we averaged the accuracy across the 50 repetitions.

### Testing for independent decoding

To compare the decoding axes between the trials and inter-trial intervals, we again trained the classifier separately for each of the three task features, but now using all the population firing rate vectors of a session, first for the trials {**r**_*t*_(1),…, **r**_*t*_(*T*)} and then for the intertrial intervals {**r**_*I*_(1),…,**r**_*I*_(*T*)}. For a given feature *f*, we then computed the angle, *θ_f_*, between the resulting vectors of decoding weights for the trials, *w_t_*(*f*), and intertrial intervals, *w_I_*(*f*), as: 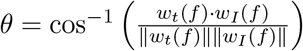. Similarly, to assess how the different features were simultaneously encoded during a task phase, we computed the angles between the decoding vectors of two features *f*_1_ and *f*_2_ within a trial, or within an inter-trial interval.

We evaluated the degree of independence between the trial and inter-trial interval decoding by attempting to cross-decode a task feature in one phase from the activity in the other. For a given task feature, we took the above classifier trained on all trials of a session, and tested its decoding on all inter-trial intervals of the same session, with performance reported as the percentage of correctly labelled inter-trial intervals. We also tested this in reverse, decoding the feature in the trials from the classifier trained to decode the same feature from all the inter-trial intervals. To check we were not overfitting when using a decoder trained on all *T* phases, we further tested cross-decoding using leave-one-out, by leaving out the *i*th trial-interval pair, training on *N* – 1 trials, and predicting the *i*th inter-trial interval (and vice-versa for training on inter-trial intervals and testing on trials). Performance for leave-one-out cross-decoding was reported as the percentage of correctly labelled held-out trials (or inter-trial intervals) over all trials of the session.

### Behavioural analysis

To check whether our decoding results depended on potentially different behaviours or task demands, we divided the sessions in two different ways, by rule type and by learning type. For the rule type, we grouped sessions by whether the target rule was a direction-based rule (so putatively egocentric) or a cue-based rule (so putatively allocentric).

To group by learning type, we identified learning sessions according to the original study’s criteria (Peyrache et al., 2009) of a session with three consecutive correct trials followed by a performance of at least 80% correct. The first of the three correct trials was the learning trial. Only ten sessions satisfied these criteria. All sessions that did not meet these criteria was labelled “Other”.

We quantified performance in learning sessions by fitting a piecewise linear regression model to the cumulative reward curve, using robust regression to fit lines before and after the learning trial. The slopes of the two lines gave us the rate of reward accumulation before (*r_before_*) and after (*r_after_*) the learning trial (Figure 1b). We quantified performance on all other sessions in a similar way (Figure 1c-d). For the 8 rule-change sessions, we considered the slopes of the regression lines before and after the rule-switch trial. For all remaining sessions, we looked for any performance change by fitting the piecewise linear regression model to each trial in turn (allowing a minimum of 5 trials before and after each tested trial). We then found the trial at which the increase in slope (*r_after_* – *r_before_*) was maximised, indicating the point of steepest inflection in the cumulative reward curve.

### Reactivation of task-feature representations in sleep

In order to quantify the reactivation of waking activity in pre- and post-session sleep, we used the population firing rate vectors computed for the decoder {**r**(1),…,**r**(*T*)}. We considered here the average population vector for a feature in each session, computed across all the trials for each feature. For example, we quantified the average population firing rate vector for all the right choice trials, and separately for all the left choice trials. This vector thus represented the region in the activity subspace (Figure 2) occupied by that particular feature in the trial or the inter-trial interval.

**Figure 2.**
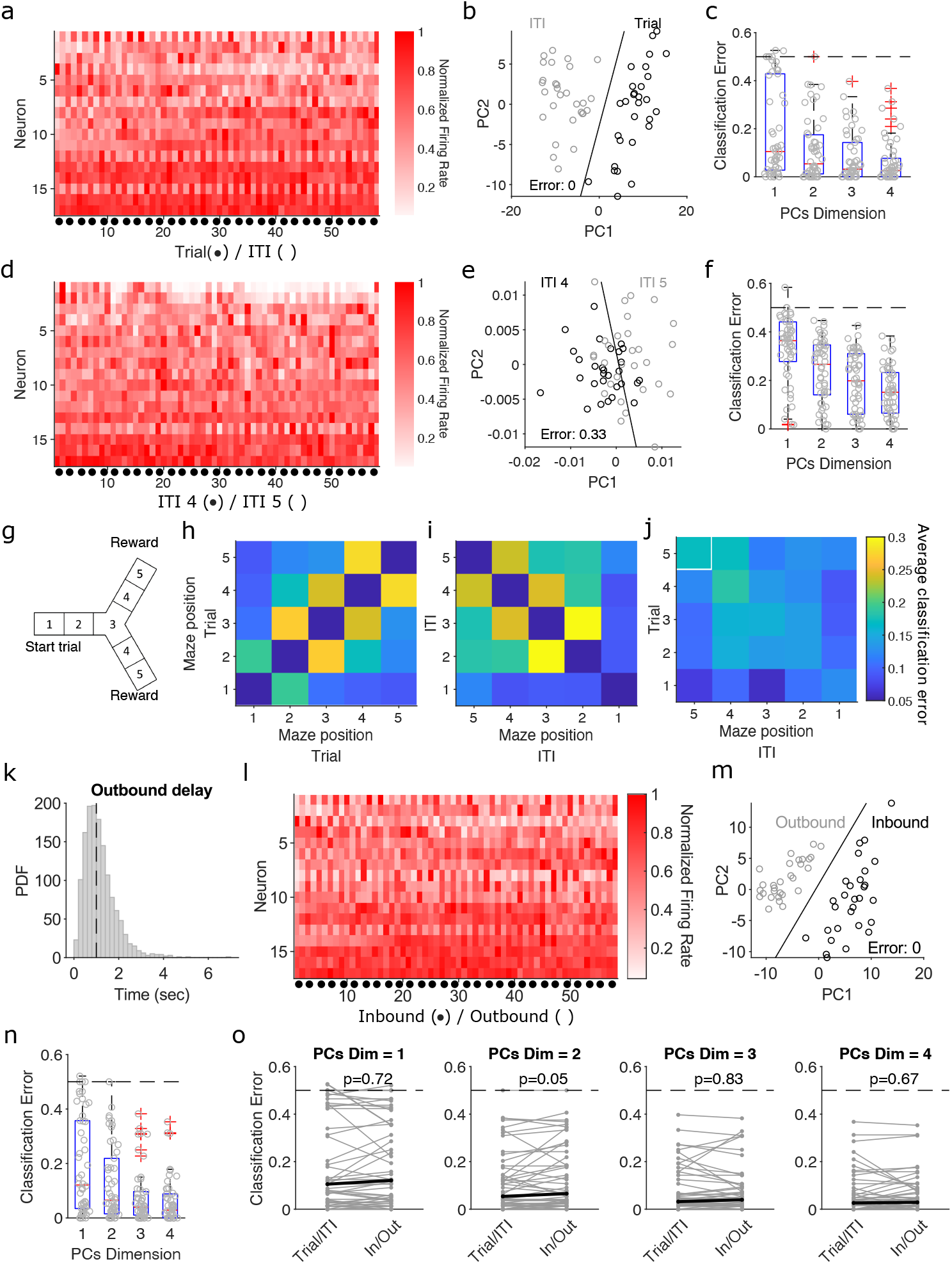
Population activity is easily separable between trials and inter-trial intervals. (**a**) Population activity vectors for the trials (•) and following inter-trial intervals (ITI) of one session with 54 trial-ITI pairs. The heat-map shows the normalized firing rate for each neuron. (**b**) Projection of that session’s population activity vectors on to two dimensions shows a complete separation of trial and inter-trial interval activity. The black line is the linear separation found by the classifier. PC: principal component. (**c**) Summary of classification error over all sessions, as a function of the number of dimensions. Each grey dot is the error for one session at that number of projecting dimensions. Dashed line gives chance performance. Boxplots show medians (red line), interquartile ranges (blue box), and outliers (red pluses). (**d-f**) Same as panels (a-c), but comparing population activity vectors for maze sections 4 and 5 in the inter-trial interval (ITI). (**g**) Schematic of maze sections. (**h**) The average classification error in the separation of population activity vectors between each pair of maze sections within trials. Classification error is for projections of the population activity vectors in a two dimensional space. (**i**) As for (h), within inter-trial intervals. (**j**) As for (h), for the separability of the trial and inter-trial interval population activity across maze sections. The white square indicates the arm-end position, where the transition from trial to inter-trial interval occurs. (**k**) Distribution of the delay between the start of the inter-trial interval and the start of the outbound phase, with a median delay of 1s (dashed line), and a mean of 1.14s. (**l**) Same session as (a), but divided by inbound and outbound phases. (**m**) As for (b), classifying vectors by inbound and outbound phases. (**n**) As for (c), for inbound and outbound vectors (**o**) Comparing the classification error for the population vectors divided by trial-ITI or by inbound-outbound phases. Each line is a session; black lines show medians. P values are from paired Wilcoxon signed rank tests.

We then compared this feature-specific activity vector with the firing rate vector of each 1 second time bin of slow-wave sleep pre- and post-session using Spearman’s correlation coefficient. This gave us a distribution of correlations between the feature-specific vector and the population activity vectors during pre-session slow-wave sleep, and a distribution of correlations between the feature-specific vector and the population activity vectors during post-session slow-wave sleep.

Spearman’s coefficient was chosen specifically to compare the relative activity of the neurons in the population between training and sleep epochs, and so we call this “reactivation”, not “replay”. Replay implies that specific patterns of firing from waking, such as sequences of place cells (Skaggs and McNaughton, 1996; Lee and Wilson, 2002; O’Neill et al., 2008; Denovellis et al., 2021) or sequences of neurons in an ensemble (Euston et al., 2007; Peyrache et al., 2009, 2010) reappear during sleep or quiescence. As we use it here, reactivation is assessing how well the sleep activity aligns with the two subspaces of trial and inter-trial interval activity during training – and consequently suggests whether those two are being revisited.

If a feature-specific activity vector was preferentially re-activated in post-session sleep, then we would expect the distribution of the correlation coefficients between a feature and post-session slow-wave sleep to be right-shifted compared to the distribution of the correlation coefficients between the same feature and pre-session slow-wave sleep. We quantified this shift by measuring the difference in the medians (*M_post_* – *M_pre_*) between the two distributions of correlation coefficients. If the difference was positive then we had a higher correlation of the feature-specific vector with the activity in post-session slow-wave sleep than with the activity in pre-session slow-wave sleep. If negative, then the feature-specific vector was more similar to the pre-session slow-wave sleep population activity.

To control for different time scales of reactivation in sleep we repeated the same procedure changing the time bin in the slow-wave sleep pre- and post-session. Bin sizes from 100ms to 10s were chosen to range below and above the mean length of a trial (about 6.5 seconds).

### Statistics

Quoted measurement values are mean *x* and SEM. Differences between two paired distributions were assessed using a paired Wilcoxon signed-rank test; differences from zero were assessed using a Wilcoxon signed-rank test. In Figures 3–7 we report where P-values for these tests exceeded three alpha levels (0.05, 0.01, 0.005). Differences between distributions were assessed using the Kolomogorov-Smirnov test. Throughout, we have *n* = 49 sessions; in some analyses we subdivide these into rule types (*n* = 15 direction-rule sessions and *n* = 34 cue-rule sessions), or learning types (*n* = 10 learning sessions and *n* = 39 Other sessions). In Figure 3 we break down the decoding results by each rat, giving *n* = 14, 14, 11, 14 sessions for rats 1-4, respectively.

**Figure 3.**
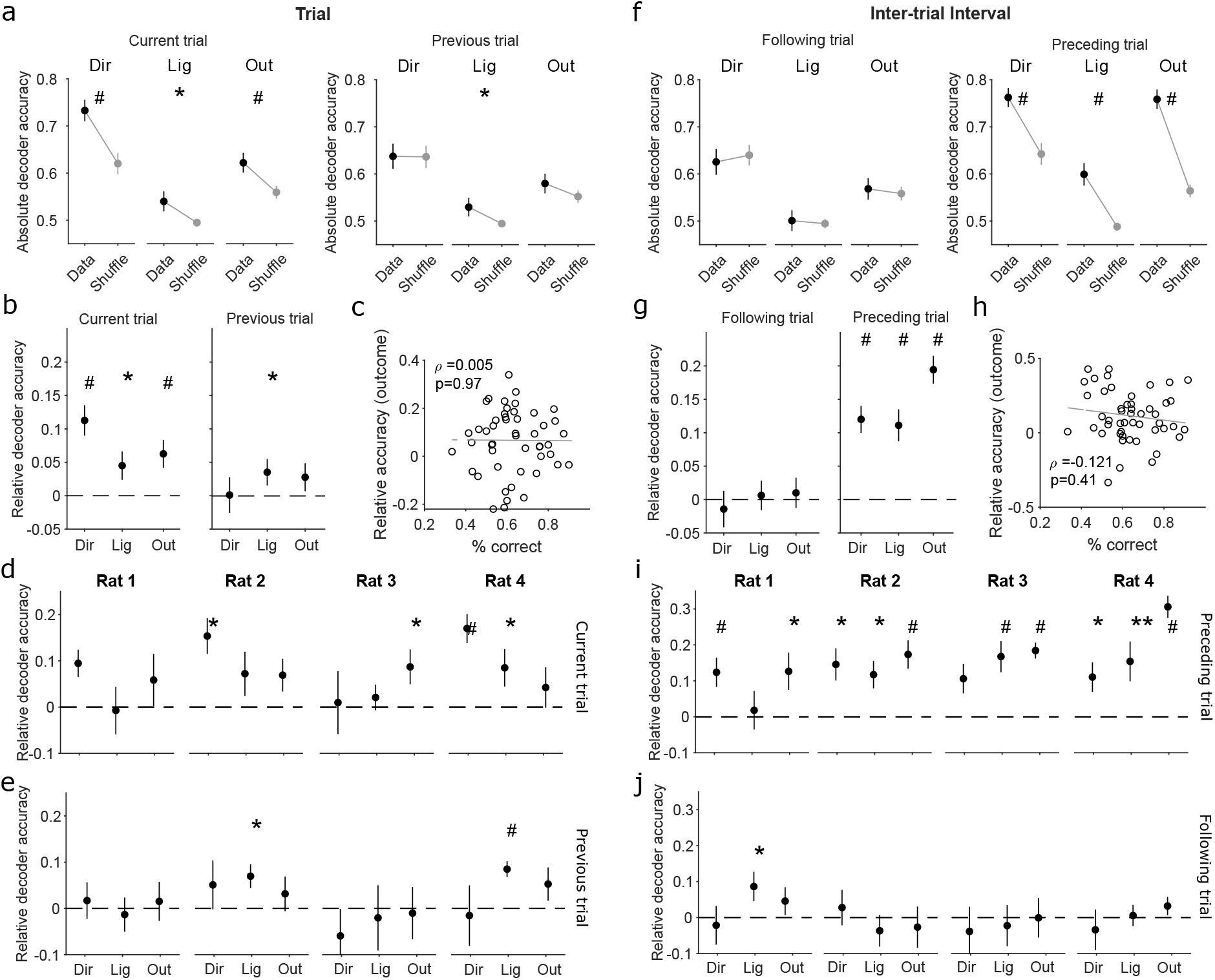
Decoding of different task states in the trials and inter-trial intervals. (**a**) Accuracy of decoding task features from population activity during trials. In black we plot the accuracy of decoding the choice of arm direction (Dir), light position (Lig), and outcome (Out) for the current trial (left panel), and the previous trial (right panel). In grey we plot the decoding accuracy of shuffled labels across trials. Differences assessed using a paired Wilcoxon signed rank test: * *p* < 0.05; ** *p* < 0.01; # *p* < 0.005. Symbols plot means ± SEM across 49 sessions. (**b**) as for panel (a), plotted as relative decoding accuracy: the difference between the decoding accuracy of the data and of the mean of the shuffled data in that session. Here and all further panels, p-values are given for a Wilcoxon signed rank test against zero median. (**c**) No correlation between session performance and the accuracy of decoding a trial’s upcoming outcome. (**d-e**) Per-subject breakdown of the decoding results in panel b. (**f**) as for panel (a), but for population activity during the inter-trial intervals of each session, decoding the features of the following (left panel) and preceding (right panel) trials. (**g**) as for panel (b), for decoding during the inter-trial interval. (**h**) as for panel (c), for decoding the outcome preceding the inter-trial interval. (**i-j**) Per-subject breakdown of the decoding results in panel g.

**Figure 4.**
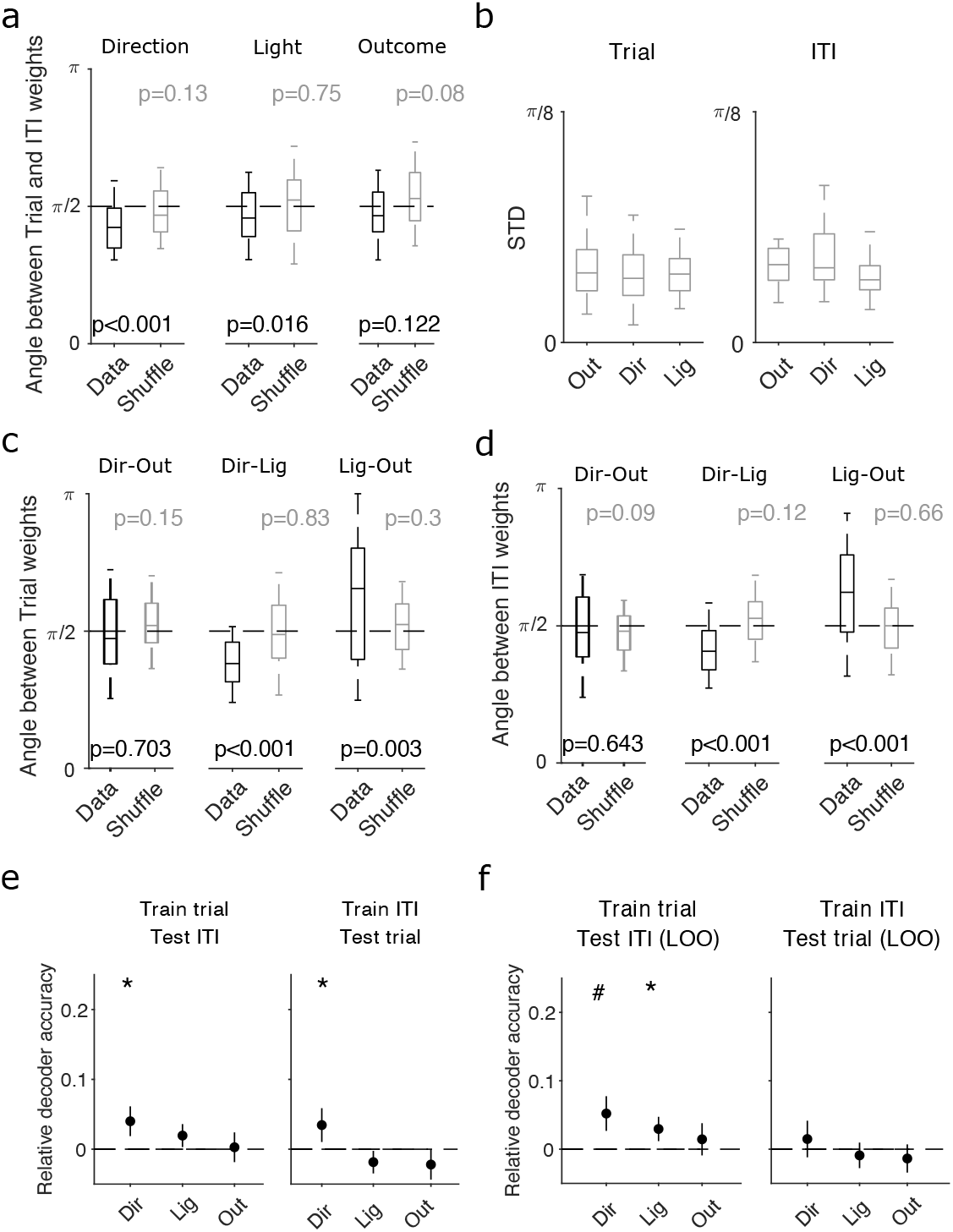
Independent decoding of task features between trials and inter-trial intervals. (**a**) Angles between the decoding weight vectors for trials and inter-trial intervals, for each task feature. For reference, we also compute the angle between trial and inter-trial interval decoding vectors obtained by training on shuffled label data (grey). Boxplots show median (line), inter-quartile range (box), and 95% interval (tails) over all sessions. P-values are from Wilcoxon ranksum tests for the difference from *π*/2. (**b**) Variation of angles between decoding axes within the same task phase. We plot here the average standard deviation of the angles between decoding axes for the same feature in either the trials (left) or inter-trial intervals (right) of a session. Within-session decoding axes for each features were taken from the classifiers fitted to each training subset of trials or inter-trial intervals during the cross-validation. (**c**) As for (a), but comparing the decoding weight vectors between features, within trials. (**d**) As for (c), within inter-trial intervals. (**e**) Cross-decoding performance for each task feature of the current trial, using the decoding vectors in panel (a). Left: performance when the decoder was trained on activity during trials and tested on activity in the inter-trial intervals. Black dashed line shows the chance level obtained training the classifier on shuffled labels for the trials and testing on inter-trial intervals given the same shuffled labels. Right: performance when the decoder was trained on activity from the inter-trial intervals, and tested on activity in the trials. * *p* < 0.05; ** *p* < 0.01; # *p* < 0.005 (**f**) As for (e), using leave-one-out cross-decoding.

**Figure 5.**
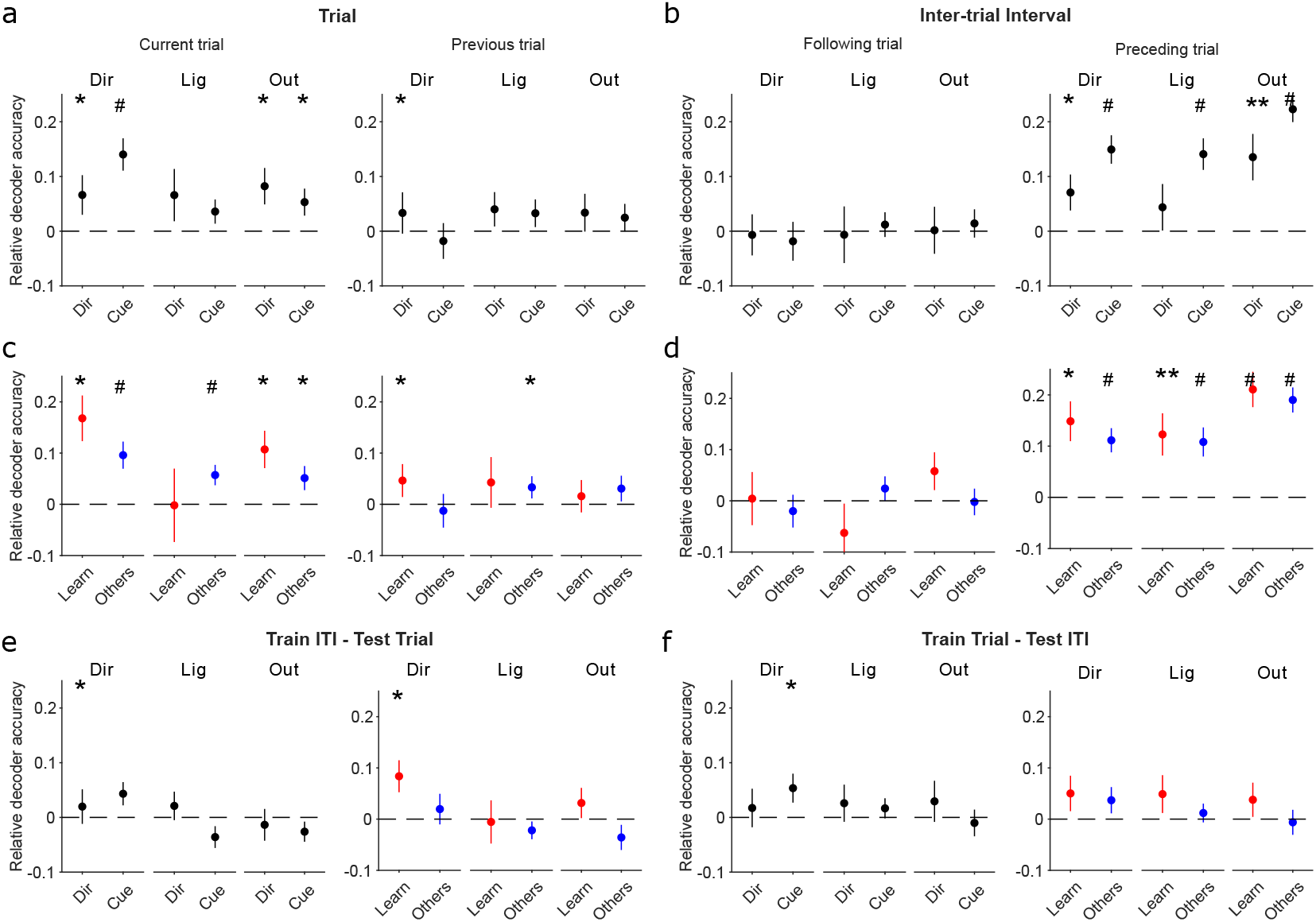
Breakdown of decoding and cross-decoding by session type. (**a**) Breakdown of the trial decoding results in Figure 3b by the rule type of each session (15 direction rule sessions; 34 cue rule sessions). * *p* < 0.05; ** *p* < 0.01; # *p* < 0.005 (**b**) As for panel (a), breakdown of the inter-trial interval decoding results by the rule-type of each session. (**c**) Breakdown of the trial decoding results in Figure 3b by whether a rule was learnt in a session or not (10 identified learning sessions; 39 other sessions). (**d**) As for panel (c), breakdown of the inter-trial interval decoding results by learning and other sessions. (**e-f**) Breakdown of the cross-decoding results in Figure 4d by rule-type (left panel) and performance-type (right panel) for each cross-decoding direction.

**Figure 6.**
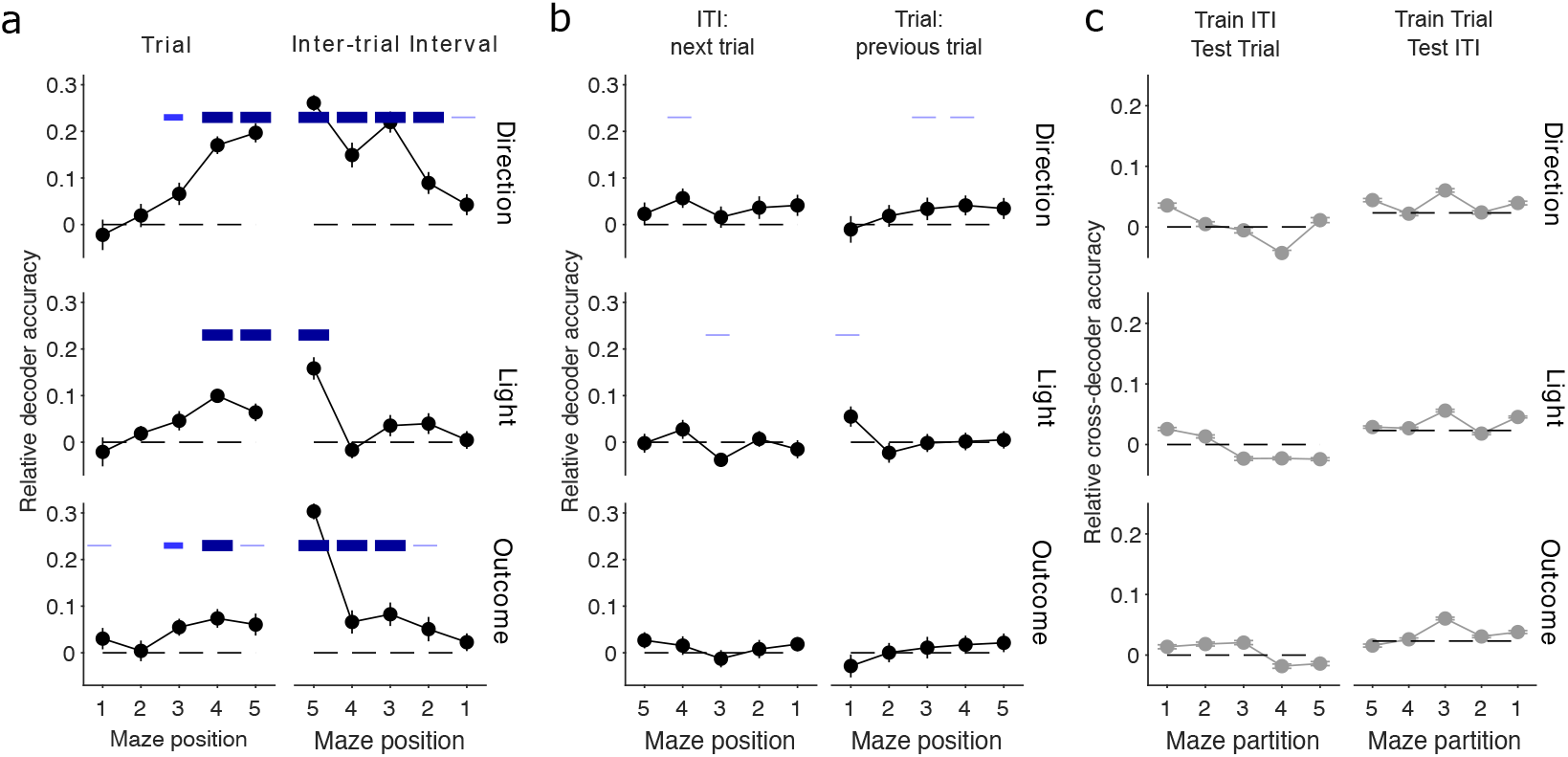
Population vector decoding across the maze. (**a**) Decoder performance at each maze location. We plot here decoding at consecutive locations across the trial and following inter-trial interval (ITI), with the decoding target for each location being the task feature in the current trial (so decoding the preceding trial during the inter-trial interval). Blue bars give significance values of: *p* < 0.05 thin blue line; *p* < 0.01 medium thickness blue line; *p* < 0.005, thick blue line, for Wilcoxon sign rank test for difference from zero. Symbols plot means ± SEM across *n* = 49 sessions. (**b**) As per (a), for: left, decoding the features of the following trial from the current inter-trial interval; right: decoding the features of the previous trial from the current trial. (**c**) Cross-decoding performance at each maze location is at chance.

**Figure 7.**
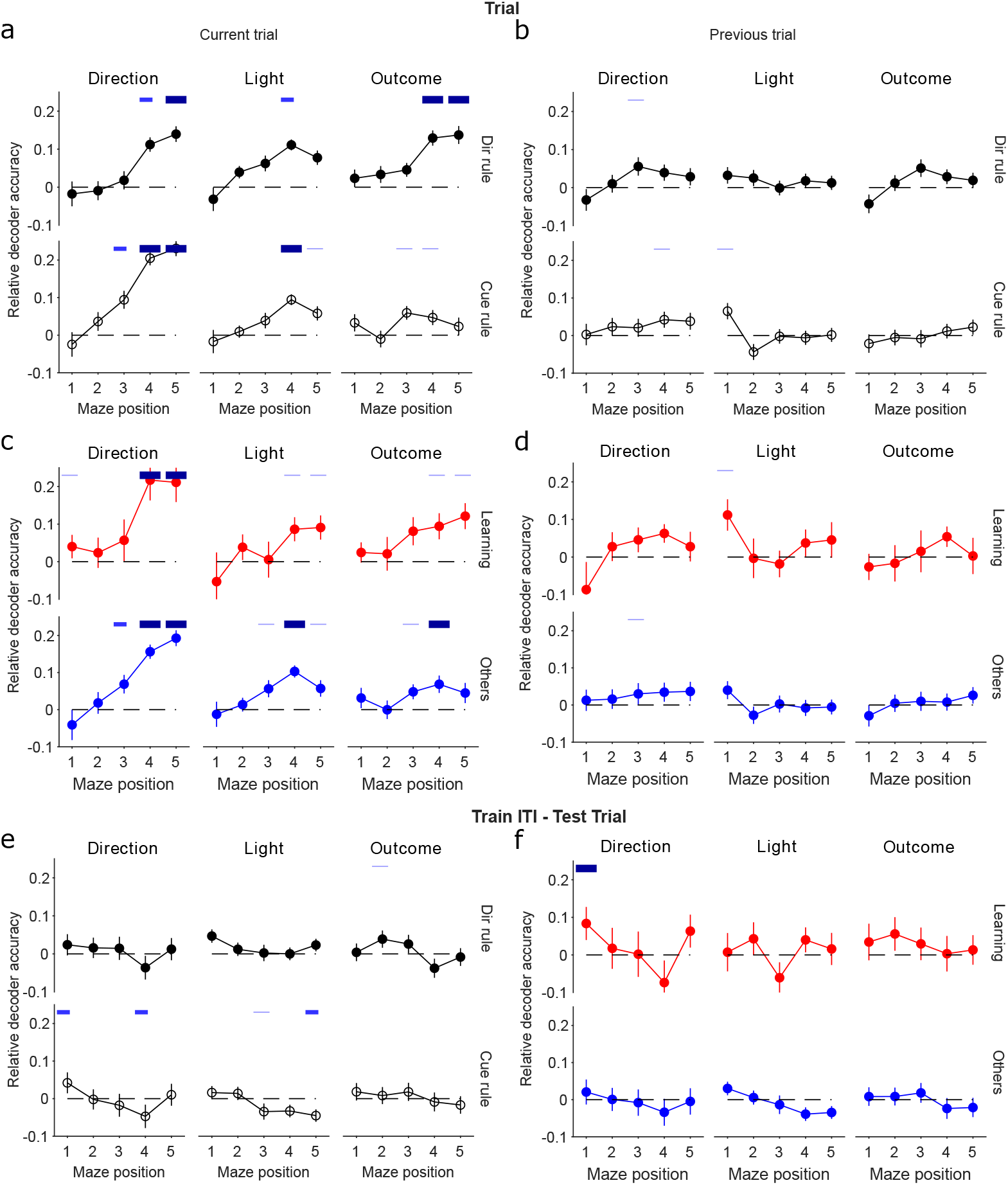
Robustness of location-dependent decoding in trials across session types. (**a**)-(**b**) Breakdown of decoding performance in Figure 6a, according to the type of rule in each session: Direction (15 sessions) or cued (34 sessions). (**c**)-(**d**) Breakdown of decoding performance in Figure 6a, according to whether the session contained putative rule learning (10 sessions, red symbols and lines) or not (39 sessions, blue symbols and lines). (**e**)-(**f**) Breakdown of cross-decoding performance at each maze location (Figure 6c) by rule-type (e) and performance-type (f). Similar results were obtained for cross-decoding inter-trial intervals from trials (not shown).

### Data and code availability

The spike-train and behavioural data that support the findings of this study are available at CRCNS.org (DOI: 10.6080/K0KH0KH5) (Peyrache et al., 2018).

## Results

We analyse here data from rats learning rules in a Y maze, who had tetrodes implanted in mPfC before the first session of training. Across sessions, animals were asked to learn one of 4 rules, which were given in sequence (go to the right arm, go to the lit arm, go to the left arm, go to the dark arm). Rules were switched after 10 correct choices (or 11 out of 12). The animal self-initiated each trial by running along the central stem of the Y maze and choosing one of the arms (Figure 1a). The trial finished at the arm’s end, and reward delivered if the chosen arm matched the current rule being acquired. During the subsequent inter-trial interval the rat made a self-paced return to the start of the central arm to initiate the next trial. Trials were 6.5 ± 0.5 seconds on average; intertrial intervals were 55.6 ± 1.1 seconds. Throughout, population activity was recorded in the prelimbic and infralimbic cortex (Figure 1e), which we shall term medial prefrontal cortex (mPfC) here (Laubach et al., 2018, propose that these regions are equivalent to the anterior cingulate cortex in primates).

Neural activity statistics differed between trials and inter-trial intervals. Neurons that had the highest firing rates in a trial tended to have lower firing rates in the following inter-trial interval (Figure 1f). However, the vector of rates across the population was not strongly correlated between the trial and following inter-trial interval (Figure 1g,h), meaning that changes in firing rates between the two phases of the task were not systematically in one direction. This low correlation of population activity between the trial and inter-trial interval is also consistent with a change in representation, as we now report.

### Separable subspaces of population activity between trials and inter-trial intervals

We first asked if population activity occupied different subspaces between consecutive pairs of trials and inter-trial intervals, as a basis for representing these as two different states of the world in the same population. To do so, we projected all population activity vectors of a session (Fig 2a) into a low dimensional space (Fig 2b), and then quantified how easily we could separate them into trials and inter-trial intervals. Using just one dimension for the projection was sufficient for near-perfect separation in many sessions; using two was sufficient for above-chance performance in all sessions (Fig 2c). Population activity thus occupied a different low-dimensional subspace between the trials and the inter-trial intervals.

This was not true within each phase (Fig 2d-f): when we divided the maze into sections (Fig 2g), the population activity at nearby maze positions was not easily separable within trials (Fig 2h) or within inter-trial intervals (Fig 2i), with the notable exception of position 1 – the return to the starting position - in the inter-trial intervals. By contrast, population activity vectors at one position in the trial and another in the inter-trial interval were easily separable between every pair of positions (Fig 2j). Thus, while of course population activity changed across maze positions (Fig 2h-i), those changes were smaller and continuous within each phase of the task, but larger and discontinuous between them, as they moved into a different subspace of activity.

A detailed examination of what might be driving this move into a different subspace of population activity is beyond our scope here, but we can show there are at least two plausible causes. Aligning population activity to the trial and inter-trial interval implies this change is caused by reaching the arm-end. But a range of other salient events may be causing this shift in population activity. To begin examining possible events, we instead divided the task into inbound and outbound phases, where the start of the outbound phase was defined from the point where the animal’s heading direction had turned towards the start of the maze for at least 400 ms, which happened on average 1.14 seconds later than the start of the inter-trial interval (Figure 2k). Dividing the population activity accordingly (Figure 2l), we found the separability of inbound and outbound phase population vectors was excellent even in only a few dimensions (Figure 2m,n).

Indeed, population separability was equally good for both the trial-ITI and inbound-outbound separations (Figure 2o). It is thus equally plausible that the shift of subspace occupied by population activity is driven by a change in heading direction as by reaching the arm-end. As these events occur close together in time (Figure 2k), considerable further work, and likely further experiments, would be necessary to tease apart the causal mechanisms (see Discussion). For the remainder of the paper we thus continue to consider activity subspaces defined by the trial-ITI split, while being mindful that exactly when the shift occurred and what causes it is unknown.

### Different states of the same task features can be decoded from population activity

We then tested if these distinct subspaces corresponded to encoding different states of the world between the trials and inter-trial intervals. Using a linear decoder on the vector of population activity during each trial (Methods), we decoded key features of the task during the trial: the animal’s choice of arm direction in the trial, the outcome of the trial, and which arm-end was lit during the trial. We trained the same decoders using the same population vectors but with features shuffled across trials (see Methods), to define appropriate chance levels for each decoder given the unbalanced distribution of some task features, such as outcome.

We could decode all of direction choice, light position, and outcome in the current trial above chance (Figure 3a,b, left panels). In Figure 3a we plot the absolute accuracy of decoding; in Figure 3b we also plot the decoding accuracy relative to the shuffled data for each session, which, as it accounts for the different distributions of features (e.g. outcome) in each session, better shows the effect size of the decoding. Relative decoding accuracies well above zero could even be seen for each animal (Figure 3d), despite the small populations (median 10 neurons, Figure 1i), the limited numbers of trials per session (median 29 trials) available for training the decoder, and the low number (11-14) of sessions. As the outcome was not yet known during the trial, the ability to decode outcome implies anticipatory activity for outcome in mPfC neurons, as previously reported by a number of labs (see Euston et al., 2012, for review). However, we found no correlation between an animal’s performance in a session and our ability to decode the upcoming outcome (Figure 3c), suggesting this anticipatory activity is not dependent on how frequently reward was acquired. Nonetheless, while it is unclear what this anticipatory activity reflects, the ability to decode outcome was robust across all classifiers we tested (not shown; see Methods and (Maggi et al., 2018)). To test for the effects of past states on population activity in the trials, we also tried decoding the direction choice, outcome, and light position of the preceding trial, and found that decoding was at or close to chance (Figure 3a,b, right panels; panel e shows each subject). Population activity in mPfC during the trials thus depended on features in the present state of the task, and weakly or not at all on features in the past trials.

In contrast, from population activity during the inter-trial interval we could decode the direction choice, outcome, and light position of the immediately preceding trial well above chance (Figure 3f,g, right), which could even be seen for each rat despite the relatively small number (11-14) of sessions each performed (Figure 3i). One caveat is that, while the light was extinguished during the inter-trial interval, precisely when is not clear from the data we have available (see Methods): consequently, it is possible that the decoding of the light could represent in part its ongoing state. Decoding of the past outcome also did not depend on performance in the session (Figure 3h). Decoding the same features of the immediately following trial was at chance (Figure 3: f,g, left, and panel j). Thus, trial and inter-trial activity both represented distinct states of the world. Moreover, the evidence suggests trial activity represented the present and inter-trial activity predominantly represented the past.

### Independent decoding axes between the trials and inter-trial intervals

Having found evidence that a single mPfC population’s activity occupies different subspaces encoding distinct states of the world, we could now ask if and how the representations are kept distinct to downstream targets.

To compare the population coding between the trial and inter-trial interval, we determined the decoding axis of trial activity for each of the present features, and the decoding axis of inter-trial interval activity for those same features in the preceding trial (Methods). These decoding axes were close to orthogonal for all three features: the angles cluster at or close to *π*/2 (or, equivalently, their dot-product clusters at or around zero) (Fig 4a). And while the decoding axes for direction choice and light position departed from purely orthogonal, the median departure was small: 0.067*π* for direction and 0.045*π* for light position. These differences between trial and inter-trial decoding axes were also consistently and substantially larger than the differences within the same phase (Fig 4b). Thus, the state of the world in the trial and inter-trial interval can be independently decoded from the same mPfC population.

By contrast, within the trial and the inter-trial interval, pairs of decoding axes for different features were not close to orthogonal, except for the comparison between the direction and outcome axes (Figure 4c,d). This neatly demonstrates that the near-orthogonality of the decoding axes between trials and inter-trial intervals is not then a trivial consequence of the decoding axes being random vectors drawn from the same distribution, because the decoding axes of the same dimension within each phase are not orthogonal. Notably, the distributions of angles between the decoding axes for a given pair of features were preserved between the trials and the inter-trial intervals, with outcome-direction around *π*/2, light-direction centered below *π*/2, and light-outcome centred above *π*/2. Thus, while each decoding axis rotated close to orthogonal between the trial and inter-trial interval, the relationships between the feature decoding axes were preserved.

To quantify how distinct these independent axes made the decoding of the trial and inter-trial states, we cross-decoded one from the other: for each feature type, we trained the classifier on all trials of a session and tested its ability to decode the same feature from their following inter-trial intervals. We found that cross-decoding was at chance level for both outcome and light position, and significant but weak for direction (Figure 4e), consistent with the angles between their decoding axes in the trials and inter-trial intervals (Figure 4a). This result was robust to whether we trained on trials and tested on inter-trial intervals, or vice-versa. Cross-decoding was also weak or at chance if we used leave-one-out testing instead (Figure 4f), by leaving out the *i*th trial-interval pair, training on *N* – 1 trials, and predicting the *i*th inter-trial interval. Thus the near independent decoding axes (Figure 4a) indeed imply that downstream targets could independently read-out either the trial or the inter-trial state of the task from mPfC population activity by using one of the two decoding axes, and use those axes to decode the current features of that state.

### Decoding and cross-decoding are robust across types of session

We explored the extent to which this decoding depended on what occurred during each session. We first split the sessions by whether the target rule was direction-based (15 sessions) or cue-based (34 sessions). For trials, the present direction choice and outcome could still be significantly decoded for both types of rule, despite the considerable drop in power from the reduced number of sessions (Figure 5a). For inter-trial intervals, the preceding direction choice, outcome, and light position could still be decoded well above chance for both types of rule (Figure 5b).

In order to determine if learning itself affected any dependence on the task state, we then separated the sessions into two behavioural groups: putative learning sessions (*n* = 10), identified by a step-change in task performance (Methods), and the remaining sessions, called here “Other” (*n* = 39). We found decoding of task features was similar when comparing learning sessions and all Other sessions for both trials (Figure 5c) and inter-trial intervals (Figure 5d). The sole exception, of decoding the current light position during trials of Other sessions but not learning sessions, could be due either to a real effect, or to the low power for decoding from 10 learning sessions.

For completeness, we also examined the breakdown of the cross-decoding results in Figure 4d by types of session. Figure 5e,f shows that cross-decoding of most features between trial and inter-trial activity remained at chance, with again significant but weak cross-decoding of direction.

### Evolution of decoding within trials and inter-trial intervals

It is likely that the decoding of task features from mPfC activity is partly dependent on maze position (Ito et al., 2015; Spellman et al., 2015). To further examine the evolution of decoding over the trial and inter-trial interval, we again divided the maze into five equally sized sections (Fig 2g), and constructed population firing rate vectors for each position. Even though the trials averaged only around 6.5 seconds in duration, and so each position was occupied for roughly one second, we still obtained clear evidence for decoding the current trial’s direction choice, outcome, and light position across multiple contiguous locations (Figure 6a, left). The contrast between the strong decoding of the current trial’s features and the weak decoding of the previous trial’s features was even clearer across maze positions (Figure 6b, left).

This evolution means that there is contiguous decoding from the trial to the intertrial interval for all three features (Figure 6a). Despite this contiguity, the cross-decoding between the same position in the two phases was at chance (Figure 6c). In particular, cross-decoding at the arm-end (position 5) was at chance, despite these being contiguous in time. This suggests that the distinct decoding of the trial and inter-trial states of the same feature appeared immediately at the arm end – or close to it (Figure 2).

Figure 7 shows that these position-dependent decoding and cross-decoding results for trials are broadly robust to breaking them down by the type of rule or by learning behaviour. Breakdowns of position decoding by session type in the inter-trial intervals are given in (Maggi et al., 2018), their Figure 5. In particular, we note here that the decoding of the state of the light during the inter-trial interval only significantly occurs at position 5 when taken over all sessions (Figure 6a, right panels), and as these data do not specify precisely when the light was extinguished during the interval, it is unknown whether that reflects the ongoing state of the light, or the past state.

### Population representations of features re-activate in sleep

That the population activity occupies linearly separable subspaces between the trial and inter-trial intervals (or the inbound and outbound phases) strongly suggests that the mPfC populations can be driven to either one or the other by upstream inputs. In turn, this implies that the representations of these two world states were independently addressable. To explore this question further, we turned to activity of the same populations during sleep.

Prior reports showed that patterns of mPfC population activity during training are preferentially repeated in post-training slow-wave sleep (Euston et al., 2007; Peyrache et al., 2009; Singh et al., 2019), consistent with a role in memory consolidation. However, these analyses looked only at specific templates or the re-appearance of correlations between neurons, so it is unknown what task states these repeated patterns represented. Thus, we took advantage of the fact that our mPfC populations were also recorded during both pre- and post-training sleep to ask if their activity during sleep was specifically driven to either or both of the activity subspaces occupied by the population during the trials and inter-trial intervals.

We first tested whether population activity representing features in the trials reactivated during slow-wave sleep. For each feature of the task happening in the present (e.g choosing the left arm), we created the mean vector of population activity specific to that feature during a session’s trials. This average population vector thus represented the region of the activity subspace (Figure 2) occupied during trials with that feature. To seek reactivation of this region of the subspace in slow-wave sleep, we computed population firing rate vectors in pre- and post-training slow-wave sleep in time bins of 1 second duration, and correlated each sleep vector with the feature-specific trial vector (Figure 8a). We thus obtained a distribution of correlations between the trial-vector and all pre-training sleep vectors, and a similar distribution between the trial-vector and all post-training sleep vectors. Greater correlation with post-training sleep activity would then be evidence of preferential reactivation of feature-specific activity in post-training sleep.

**Figure 8.**
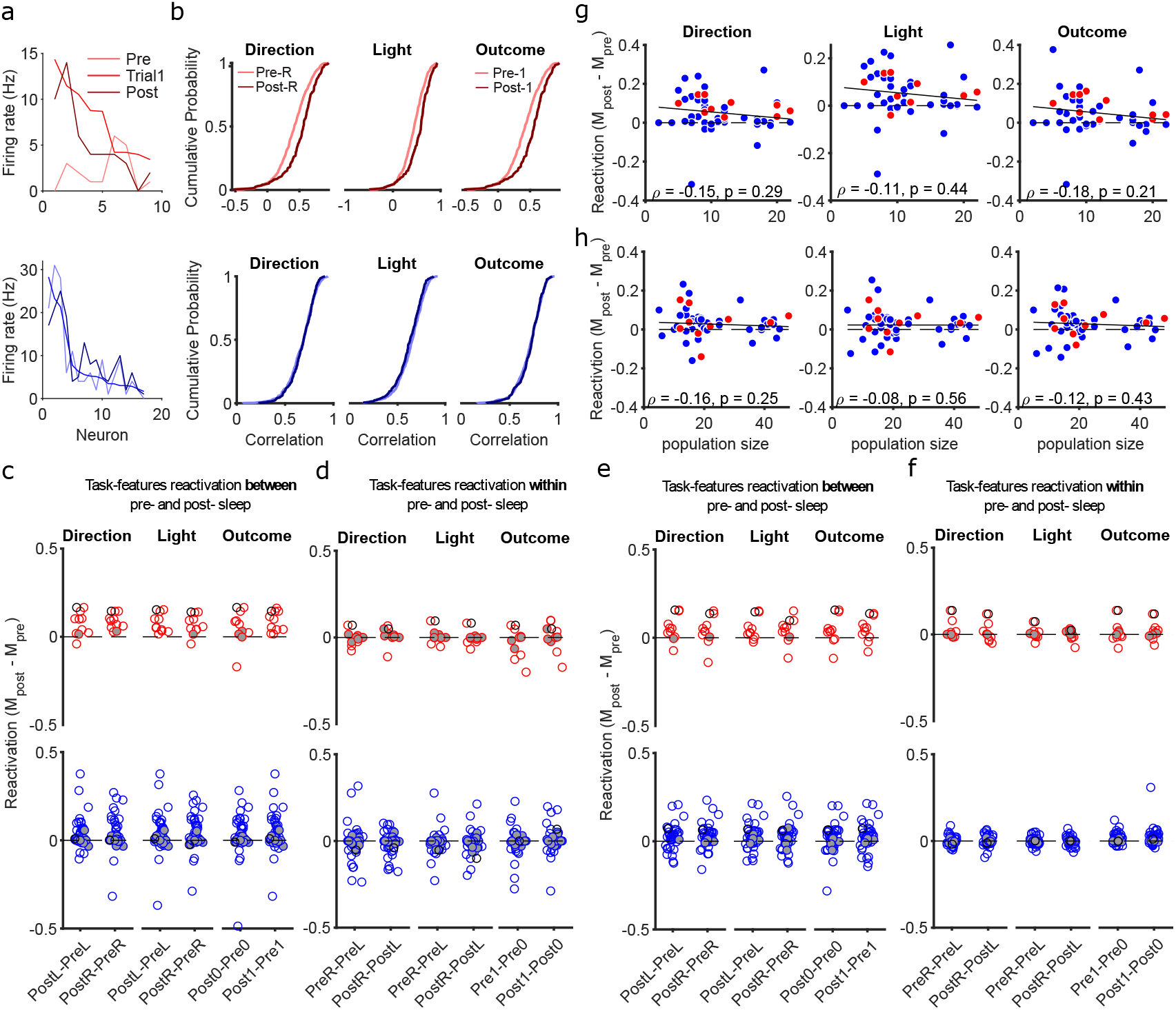
Reactivation of population subspaces in post-training sleep. (**a**) Example population activity vectors. Upper panel: from one learning session, we plot the average firing rate vector for correct trials (Trial1). For comparison, we also plot examples of firing rate vectors from pre- and post-training slow-wave sleep (1s bins). Neurons are ranked in order of their firing rates in the trial vector. Lower panel: as for the upper panel, for an example session not classified as learning. (**b**) Example distributions of Spearman’s rank correlations between trial and sleep population activity. Upper panels: for the same learning session as panel (a), we plot the distributions of correlations between each vector of feature-specific trial activity and the population activity vectors in pre- and post-training slow-wave sleep. Lower panels: as for the upper panels, for the example non-learning session in panel (a). R: right arm; 1: rewarded trial. (**c**) Summary of reactivations of trial activity across all sessions. For each task feature, we plot the difference between the medians of the pre- and post-training correlation distributions. A difference greater than zero indicates greater correlation between trials and post-training sleep. Each symbol is a session. Empty symbols are sessions with significantly different correlation distributions at *p* < 0.05 (Kolmogorov-Smirnov test). Grey filled symbols are not significantly different. One black circle for learning and one for non-learning sessions identify the two example sessions in panels (a) and (b). (**d**) As for panel c, but plotting the median differences between distributions for paired task features within the same sleep epoch. For example, in the left-most column, we plot the difference between the correlations with pre-session sleep activity for right-choice and left-choice specific trial vectors (PreR - PreL). (**e** - **f**) As for panels c-d, for inter-trial interval activity. (**g**) Relationship between population size and the strength of preferential reactivation of trial activity. Symbols show sessions: red are learning, blue are Other sessions. Solid line gives the best-fit linear regression; text gives Spearman’s rank correlation. (**h**) As for panel g, for inter-trial interval activity.

We examined reactivation separately between learning and Other sessions, seeking consistency with previous reports that reactivation of waking population activity in mPfC most clearly occurs immediately after rule acquisition (Peyrache et al., 2009; Singh et al., 2019). Figure 8b (upper panels) shows an example of a learning session with preferential reactivation. For all trial features, the distribution of correlations between the trial and post-training sleep population activity is right-shifted from the distribution for pre-training sleep. For example, the population activity vector for choosing the right arm is more correlated with activity vectors in post-training (Post-R) than pre-training (Pre-R) sleep.

Such post-training reactivation was not inevitable. In Figure 8b (lower panels), we plot another example in which the trial-activity vector equally correlates with population activity in pre- and post-training sleep. Even though specific pairs of features (such as the left and right light positions) differed in their overall correlation between sleep and trial activity, no feature shows preferential reactivation in post-training sleep.

These examples were recapitulated across the data (Figure 8c). In learning sessions, feature-specific activity vectors were consistently more correlated with activity in post-than pre-training sleep. By contrast, the Other sessions showed less consistent preferential reactivation of any feature-specific activity vector in post-training sleep. As a control for statistical artefacts in our reactivation analysis, we looked for differences in reactivation between paired features (e.g. left versus right arm choice) within the same sleep epoch and found these all centre on zero (Figure 8d). Thus, population representations of present task features in the trials were preferentially reactivated in post-training sleep, and this most consistently occurred after a learning session.

We repeated the same analyses using feature-specific population vectors from the inter-trial interval activity and also found evidence of preferential reactivation in some sessions (Figure 8e-f). However, in contrast to trial activity, there was no consistent preferential reactivation of inter-trial interval activity after a learning session.

Neither the preferential reactivation of trial nor inter-trial activity was explained by significantly higher correlations between waking and sleep activity vectors from smaller populations (Figure 8g-h).

As our measure of reactivation is asking if and when the mPfC population’s activity revisits the trial and/or inter-trial activity subspaces, it could do so on a range of time-scales. These patterns of preferential reactivation were consistent across a range of binsizes used to construct the activity vectors during sleep (Figure 9). Notably, across these time-scales, trial activity showed two independent properties from inter-trial interval activity: consistent preferential reactivation after learning sessions, and that preferential reactivation in those sessions was stronger at smaller binsizes. These results are consistent with trial and inter-trial activity subspaces being independently addressable; we thus sought further evidence of their independence.

**Figure 9.**
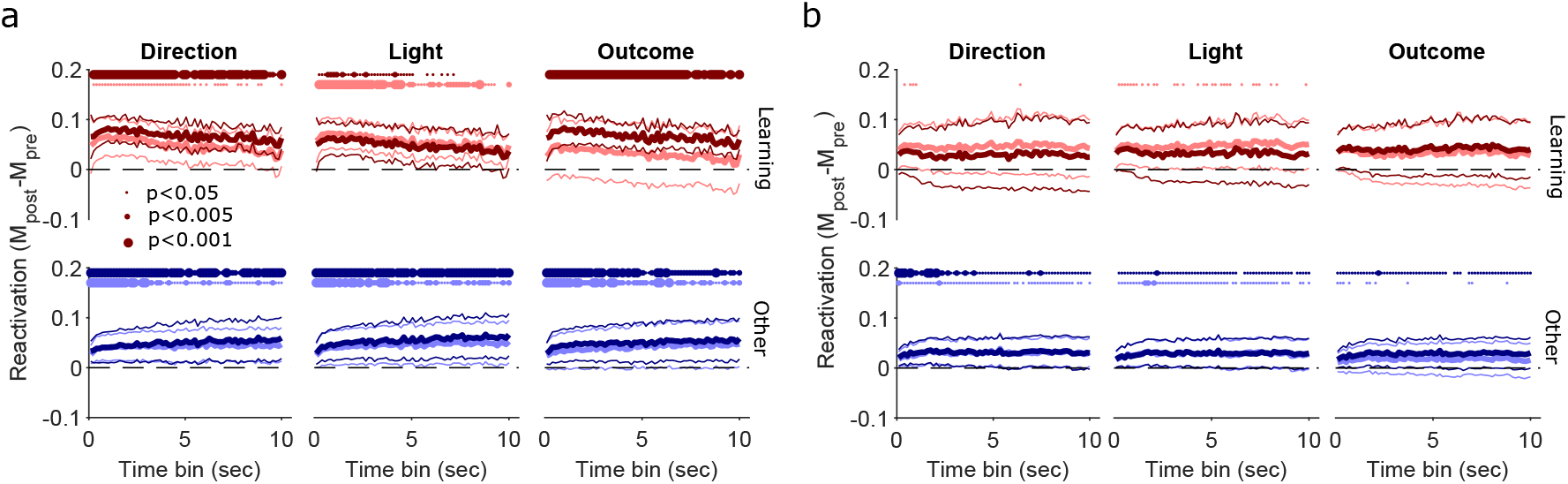
Robust preferential reactivation across time-scales of sleep activity. (**a**) Preferential reactivation of trial activity across time-scales. At each time bin used to construct population activity vectors in sleep, we plot the distribution over sessions of the median differences between pre- and post-training correlation distributions, for learning (top), and other (bottom) sessions. Distributions are plotted as the mean (thick lines) ± 2 SEM (thin lines); at the 1s bin, these summarise the distributions shown in full in Figure 8c. Each panel plots two distributions, one per pair of features: lighter colours indicate left or error trials (L or 0); while darker colours indicate right or correct trials (R or 1). Time bins range from 100 ms to 10 s, tested every 150 ms. Dotted lines at the top of each panel indicate bins with reactivation significantly above zero (Wilcoxon sign rank test, *p* < 0.05 thin dot; *p* < 0.005 middle size dot; *p* < 0.001 thicker dots; *N* = 10 learning, *N* = 39 Other sessions). (**b**) As for panel a, for inter-trial activity.

### Independent properties of trial and inter-trial activity reactivation in sleep

We asked whether the amount of reactivation of population activity differed between trial and inter-trial activity. The reactivation of trial population activity was strongly correlated between pre- and post-session sleep (Figure 10a), but the reactivation of inter-trial interval activity was less correlated (Figure 10b), and this was consistent across time-scales used to construct the sleep activity vectors (Figure 10c). Thus the overall reactivation of trial and inter-trial interval activity was consistently different, again suggestive that the two subspaces of activity were independently addressable.

**Figure 10.**
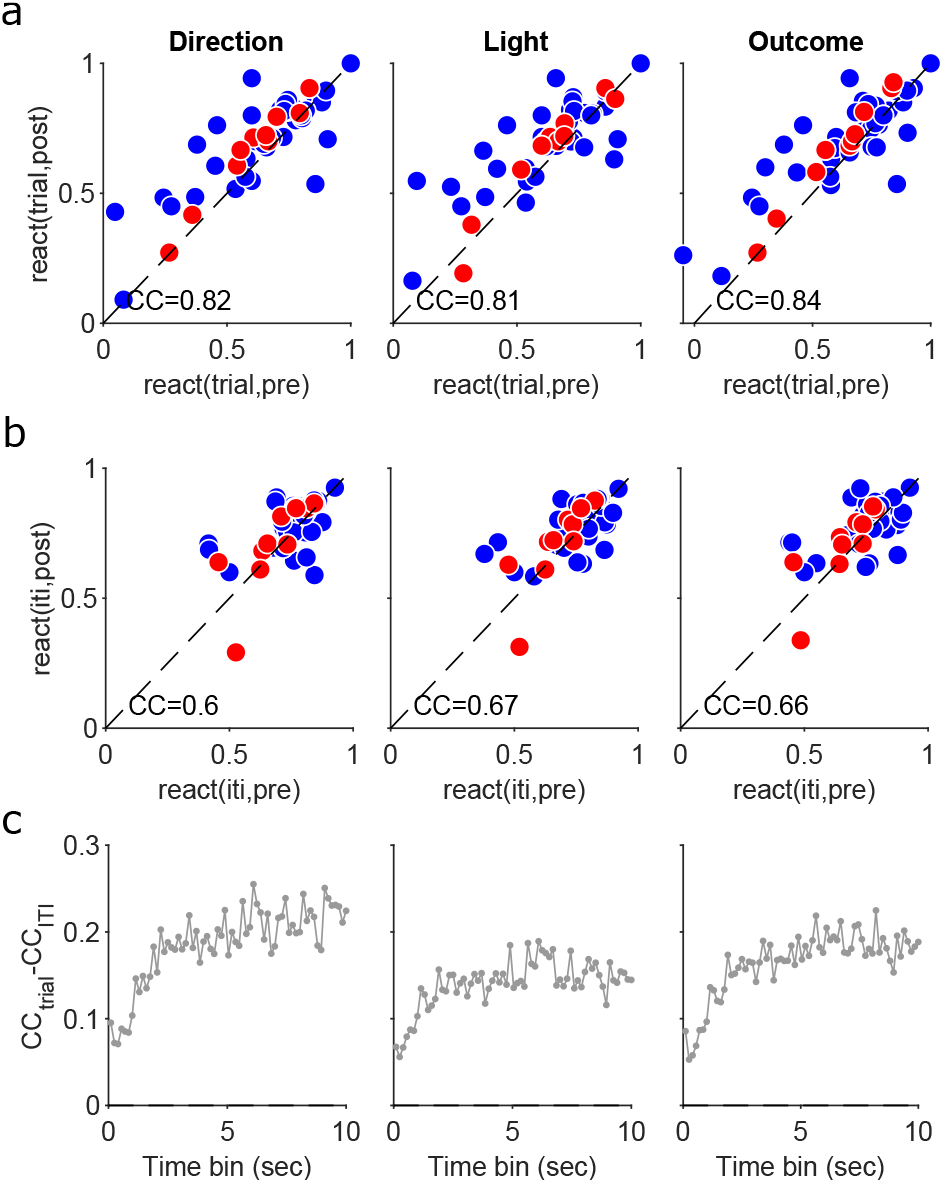
Trial activity reactivation is more correlated between pre- and post-session sleep. (**a**) Correlation between the reactivation of trial activity in pre- and post-session sleep for each task feature. Reactivation is as defined in Figure 8a: the median rank correlation between the waking and sleep activity vectors. Note the preferential reactivation is shown by symbols falling above the dashed identity line. Sleep activity vectors constructed using 1s bins. CC: correlation coefficient. Red: learning; blue: Other. (**b**) As for panel a, for inter-trial activity. (**c**) Difference between trial and inter-trial activity reactivation correlations across binsizes used to construct sleep activity vectors.

Given the above evidence that reactivation of trial and inter-trial interval activity could be independently controlled, we further asked if they differed in how preferential reactivation correlated with behaviour. Following the differences in reactivation after learning sessions (Figure 9), we looked at the degree of learning in a session, which we quantified by the size of the change in reward rate in that session (Methods). We found preferential reactivation of trial activity correlated with the change in reward rate (Figure 11a), but preferential reactivation of inter-trial activity did not (Figure 11b). Again, this difference between trial and inter-trial activity reactivation was consistent across a wide range of time-scales used to construct the sleep activity vectors (Figure 11c-d).

**Figure 11.**
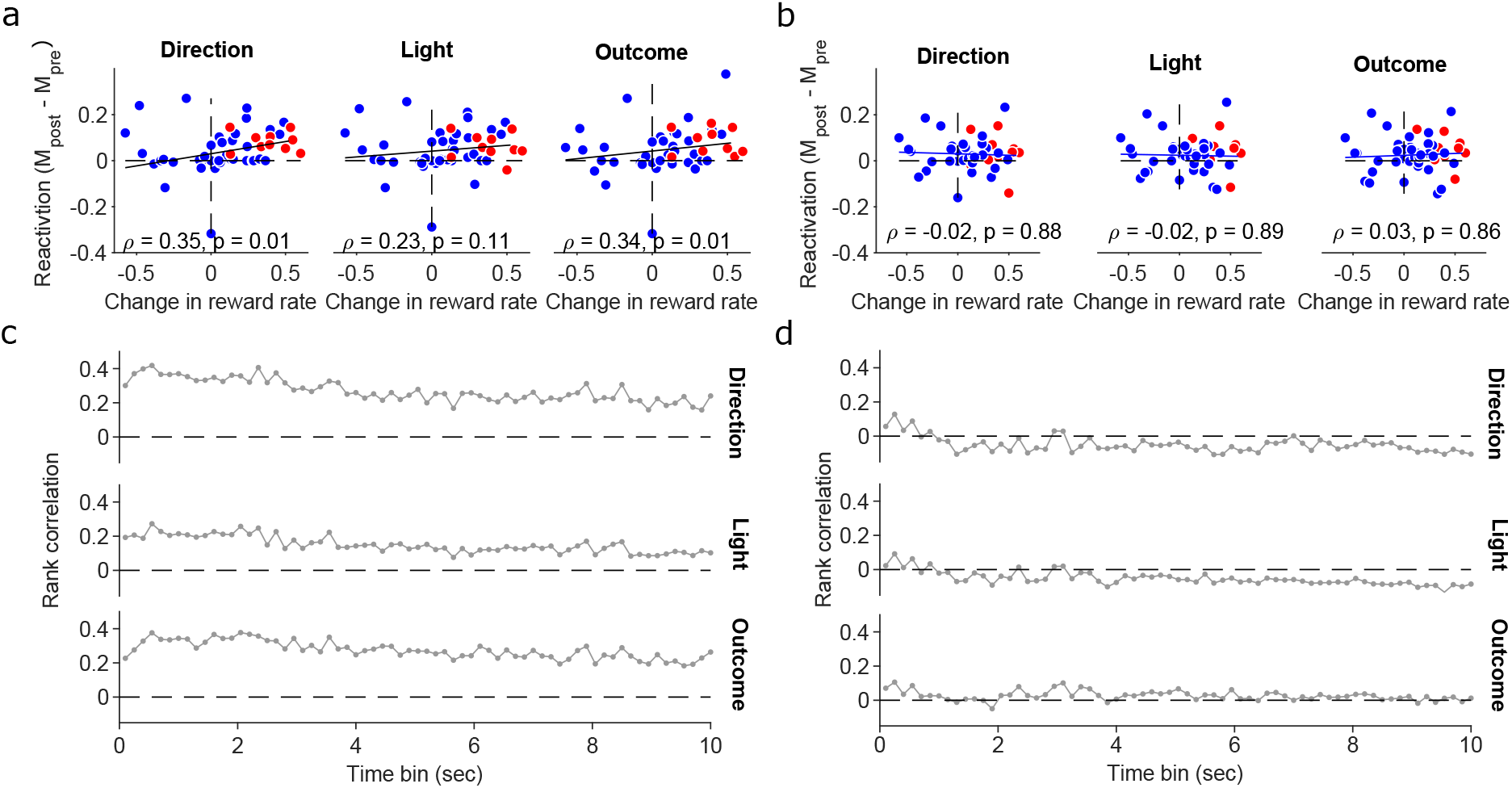
Preferential reactivation of trial activity correlates with performance. (**a**) Correlation between performance and preferential reactivation of trial activity. Sleep activity vectors constructed using 1s bins. Values are Spearman’s rank correlations. We plot here reactivation of vectors corresponding to left (direction and light) or correct; correlations for other feature-specific vectors are similar in magnitude: 0.37 (choose right), 0.35 (cue on right), 0.2 (error trials). (**b**) As for panel a, for inter-trial interval activity. Correlations for other feature-specific vectors are similar in magnitude: −0.004 (choose right), 0.02 (cue on right), −0.08 (error trials). (**c**) Correlations between performance and preferential reactivation of trial activity across binsizes used to construct sleep activity vectors, for the features in panel a. (**d**) As for panel c, for inter-trial interval activity.

## Discussion

Activity in the prefrontal cortex is known to represent different states of the world, including the immediate past or present state in a range of tasks (Baeg et al., 2003; Averbeck et al., 2006; Fujisawa et al., 2008; Rigotti et al., 2013; Sul et al., 2010; Ito et al., 2015; Hanks et al., 2015; Spellman et al., 2015; Siegel et al., 2015; Guise and Shapiro, 2017). How the representations of the different states relate to each other, and whether they co-exist in the same population of neurons, has been unclear. Consequently, it is unknown how downstream readouts of prefrontal cortex activity can distinguish activity representing different states of the world.

Here we have shown one potential solution in the medial prefrontal cortex of rats learning rules in a Y-maze, that different states are encoded in the same population in such a way that linear decoders can read-out different states of multiple features of the task. That encoding had two notable features. First, that population activity is linearly separable between the trial and inter-trial interval in as little as one dimension, so exists in different subspaces during these two phases of the task. Second, that the decoding was roughly orthogonal between the trial and inter-trial activity. These two features allow a simple solution to the interference problem.

### The interference problem

Any neural population whose activity contains information about multiple states of the world faces the problem of interference (Libby and Buschman, 2021), of how downstream populations can distinguish the activity that depends on each state, so that the sequence and causality of world events is clear. The inverse problem is how inputs to the population can selectively recall only the activity that depends on a particular state.

As we have shown here, because trial and inter-trial activity occupies different subspaces of the population activity, a downstream target using a linear decoder can distinguish the two (Semedo et al., 2019). This suggests a simple solution to the interference problem for mPfC activity, of having two downstream populations, one whose input weights from the mPfC population match the decoding axis for the trial state, and another whose input weights from the mPfC population match the decoding axis for the inter-trial state. Then the first downstream population only responds to activity representing the trial’s state, and the other only to activity representing the inter-trial’s state: and these potentially map onto the present and the past – see below.

Key here is that the decoding axes are orthogonal, or close to it, even though the population activity in mPfC is not. In Figure 12 we show this by plotting the angles between the mean activity vectors representing each feature in trials and inter-trial intervals: we see that the activity representing each feature is more closely aligned between trials and inter-trial intervals than are the corresponding decoding axes. Despite this alignment, because the activity sits in different linearly separable subspaces between trials and intertrial intervals, so the different states of the task in the trial and in the inter-trial interval are easily distinguishable by a linear decoder.

**Figure 12.**
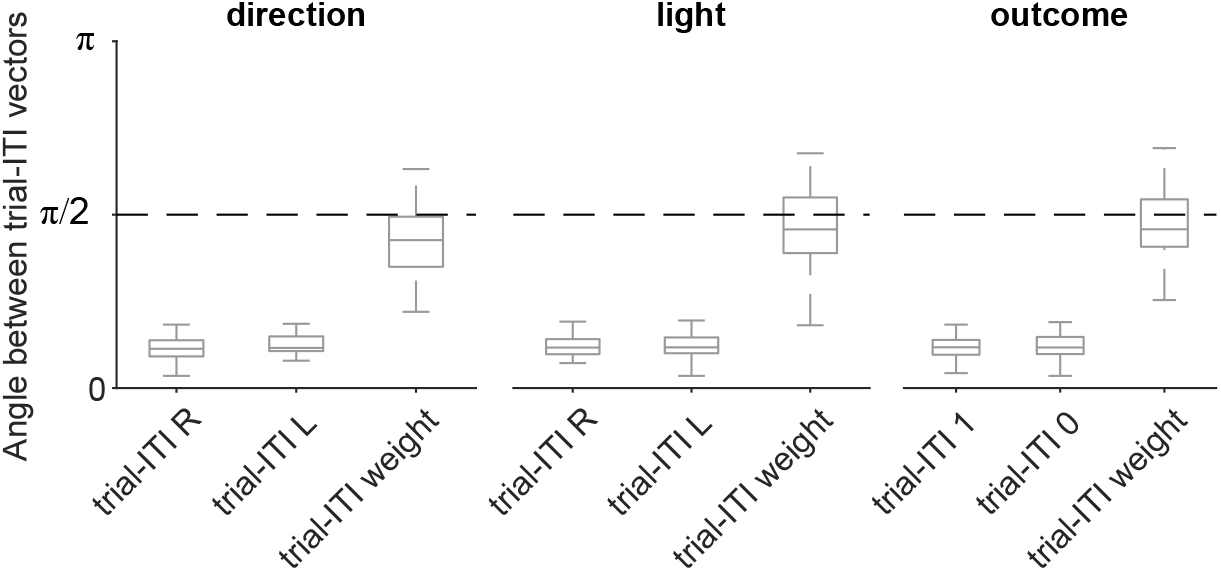
Population activity is not orthogonal, only decoding axes. For each feature (e.g. choose left), we plot the distribution of angles between the corresponding mean activity vector in trials and inter-trial intervals. We also re-plot from Figure 4a the distribution of angles between the trial and inter-trial interval decoding axes for that feature. L: left; R: right: 1: reward; 0: no reward.

We also found evidence here of a solution to the inverse problem, as the existence of different subspaces of activity between trials and inter-trial intervals means that upstream inputs could in principle separately drive the population activity to either subspace. The sequential decoding also strongly suggests that the same mPfC population can be driven into different representations by upstream inputs.

To explore this further we looked at activity of the same mPfC populations during sleep, to ask if trial and inter-trial intervals representations of the task features are reactivated differently. Trial and inter-trial activity were both preferentially reactivated in post-training slow-wave sleep, yet we found evidence that preferential reactivation of trial activity differed in four ways: the time-scales at which it occurred most strongly, that it occurred after learning sessions, that the strength of reactivation was more consistent between pre- and post-session sleep, and that it correlated with the rats’ performance in the sessions. Together, these differences between the reactivation of trial and inter-trial interval activity are consistent with upstream inputs to the mPfC population being able to separately address the representations of these states.

The consistency of preferential reactivation across broad time-scales suggests that it is the changes to the relative excitability of neurons within the mPfC population that are carried forward into sleep (Singh et al., 2019). Thus, this consistency across broad time-scales implies that whenever the population’s neurons are active, they are active together with approximately the same ordering of firing rates.

### Mixed population coding in mPfC

Our finding that small mPfC populations can sustain mixed encoding of two or more of the current trial’s direction choice, light position, and outcome is consistent with prior reports of mixed or multiplexed coding by single neurons in the prefrontal cortex (Jung et al., 1998; Horst and Laubach, 2012; Rigotti et al., 2013; Fusi et al., 2016; Aoi et al., 2020). These encodings were also position-dependent. Decoding of direction choice reliably occurred from the maze’s choice point onwards, but it is unclear whether this represents a causal role in the choice itself, or an ongoing representation of a choice being made.

Indeed, we are not claiming that the specific task features we decoded are necessarily explicitly represented in mPfC population activity. Rather, throughout we have interpreted the decoding of these features as evidence that mPfC population activity is at least representing the state of the world, similar to reinforcement learning views of PfC representations (Wang et al., 2018), because these features are a part of that state; and hence any change in one of those features, such as arm choice, would thus be a different state of the world.

Previous studies have reported that past choices modulated mPfC population activity during trials (Baeg et al., 2003; Sul et al., 2010). In contrast to the robust decoding of the present, we found weak evidence that mPfC activity during a trial depended on the light position of the previous trial, and weak evidence that it depended on the previous trial’s direction choice only during direction-based rules. Moreover, these features of the past could only be decoded at one or two locations on the maze. Thus, during trials, population activity in the prefrontal cortex had robust, sustained dependence on multiple features of the present, but at best weakly and transiently depended on one feature of the past.

Indeed, we have evidence here that the trial and inter-trial activity represent not just different task states, but respectively the present and past state of the task: trial activity decoded present but not past features; inter-trial activity decoded features of the preceding trial. The latter is consistent with well-established roles for the prefrontal cortex in shortterm memory (e.g. Funahashi et al., 1989; Machens et al., 2010; Constantinidis et al., 2018; Lundqvist et al., 2018). However, the limitations of the Y-maze task data mean we cannot rule out that the inter-trial activity also represented some features of the present during that interval - a question to be pursued further. Nonetheless, we have strong evidence that mPfC activity represents distinct states in the trials and in the inter-trial intervals.

### What could drive changes in mPfC population activity

The evolution of activity within trials and inter-trial intervals was continuous, with adjacent maze sections containing more similar population activity, yet the transition from the trial to the inter-trial interval was discontinuous, with population activity moving to a different subspace, linearly separable from the trials’. What might be driving this shift from the trial to the inter-trial interval subspace of activity, and hence its decodability?

The division into trials and inter-trial intervals or the inbound and outbound phases in Figure 2 both distinguish the two legs of the journey in the maze. During the return trip to the starting position, the change in context and direction of movement would likely change the signals available to the mPfC. It does not automatically follow though that changes in context and movement cause the observed changes in population activity in mPfC: those changes to sensory and movement information could have changed mPfC population activity so that it did not encode anything about the immediately preceding trial – in the same way, for example, that we showed the inter-trial activity encodes nothing about the immediately upcoming trial, even when the trial’s decision could be known in advance. Thus, our finding that we could still decode the state of the immediately preceding task features from inter-trial activity despite the changes in context and movement information is non-trivial. Indeed, it implies that those changes could be the drivers of the observed changes in population activity.

This suggests multiple lines of fruitful further work here. One open question is what inputs to the mPfC drive the move from one activity subspace to another. Given the switch in context and heading direction, a likely candidate is the direct input from region CA1 of the hippocampus (Jones and Wilson, 2005; Benchenane et al., 2010, 2011). Another open question is precisely when the change in activity subspace happens: we showed here preliminary results that the larger, discontinuous change in population activity could be plausibly either upon reaching the arm-end or on initiating the outbound trip back to the starting position. Another is the precise function of the representations of the trial and inter-trial interval: one possibility is they respectively reflect reward prediction and reward processing. One possibility for tackling this question is to examine how much the clean independence between the decoding of task states depends on the behavioural task. For example, tasks where the future choice of arm depends on recent history, such as double-ended T-mazes (Jones and Wilson, 2005), multi-arm sequence mazes (Poucet et al., 1991), or delayed non-match to place (Spellman et al., 2015), blur the separation of the present and the past. Comparing population-level decoding of the states in such tasks would give useful insights into when they are or are not independently coded within mPfC.

## Acknowledgments

We thank Adrien Peyrache for the data, discussions, and comments on early drafts of this manuscript, Hazem Toutounji and Martin O’Neill for comments on drafts, and the Humphries’ lab past and present (Abhinav Singh, Javier Caballero, Mat Evans, Francois Cinotti, Tomas Fiers) for discussion. This work was supported by the Medical Research Council [grant numbers MR/J008648/1, MR/P005659/1 and MR/S025944/1]. The original data collection was supported by the EU Framework (FP6) “ICEA” grant.

## Author Contributions

M.D.H and S.M. designed the analyses. S.M. analysed the data. M.D.H and S.M. wrote the manuscript.

